# CNN-based radiographic acute tibial fracture detection in the setting of open growth plates

**DOI:** 10.1101/506154

**Authors:** Zbigniew A. Starosolski, Herman Kan, Ananth V. Annapragada

## Abstract

Pediatric tibial fractures are commonly diagnosed by radiographs and constitute one of the common tasks performed by pediatric radiologists. Here, we assess the performance of a convolutional neural network for the detection of acute tibial fractures trained with a limited number of cases in skeletally immature patients. This retrospective study was performed on radiology reports manually classified as normal or tibial fracture. Classified images of orthopaedic implants, casting, and images including other pathology were excluded. The remaining cases constituted 516 studies containing 2118 radiographs. These radiographs were truncated to include a limited investigated field of view which included the distal third of the leg, inclusive of the distal physis. After exclusions, the culled dataset was randomly divided into a training set containing 784 radiographs, a validation set containing 98 radiographs, and a test set 98 radiographs. We used a modified transfer learning approach based on the Xception architecture with additional fully convoluted reasoning and drop-out layers. Of 49 fractures, two were misdiagnosed as normal. Of 49 normal exams, none were misdiagnosed. This led to model accuracy of 97.9%, sensitivity 95.9%, and specificity 100%, comparable to or better than human radiologists. In no instances were normal physes or normal developmental epiphyseal fragmentation of the tibial tuberosity or medial malleolus misclassified as a fracture. We report an efficient method to use a pre-trained network and adapt it to a medical classification task using only a small number of radiographs dedicated to precise anatomical location.

## Introduction

Subspecialty radiology training has a key role in accuracy with second opinion overreads at a pediatric specialty hospital differing from the original reads from a community, general radiologist in ∼42% of the cases (refer eakins et al.). Second reads within specialty however, exhibited substantially lower discrepancy rates, 2-4%, even when comparing resident/fellow reads and attending physician reads (refer Bisset, kung, tomich), with the largest discrepancies observed in fracture for appendicular radiographs and intracranial hemorrhage for head CT in the setting of trauma. Simultaneously, the demand for rapid turnaround and triage for pediatric orthopaedic imaging referrals after trauma has increased steadily at our institution and elsewhere. For example, pediatric orthopaedic imaging for the evaluation after trauma and non-weightbearing constitutes approximately 20% of our radiologic emergency room (ER) and urgent care volume. At our institution, one pediatric radiologist covers 3 ER (one level 1, two level 2) centers and 13 urgent care centers during the after-hours period. The need for rapid report generation and triage is limited by the speed of the radiologist interpreting the radiographs, and therefore a clear need exists for improved workflow management, in which artificial intelligence (AI) augmented interpretation has clear promise [1].

Previously tried techniques for fracture detection in pediatric patients have failed [2] due to the wide range of anatomic regions, size, projection angles, skeletal maturation, vast differences in imaging hardware, and distractions due to non-relevant objects or extraneous anatomy (eg. parent’s hand holding the child’s extremity) in recorded images. Due to recent improvements in computational hardware, machine learning has exhibited the potential to solve these problems. Convolutional Neural Networks (CNN’s) are one of the methods that show promise in this application. CNN architecture mimics neural architecture in the brain using an interconnected network of artificial neurons. These basic units perform simple mathematical operations on inputs and produce only one output. Each artificial neuron performs two consecutive operations 1) a weighted sum of inputs, and 2) passing the results of the sum through an activation function. Activation function could be linear on non-linear, it maps the input value from 0 to 1 or −1 to 1 depending on the chosen function type. Usually, activation functions have a sigmoid shape.

Artificial neurons are structured in layers. Depending on the network architecture, artificial neuronal layers are connected to functional blocks performing nonlinear operations such as pooling and n-dimensional convolutions. Repeated layers are then stacked creating a “deep” network. Interconnections between layers define distinct deep networks. The Network training/optimization process adjusts each weight to minimalize a chosen penalty function. The prediction is made (calculated) at the last network layer, based on inputs provided to network using the weights that resulted from training [3].

The accuracy of the prediction depends strongly on network architecture and data available for training. The training data should adequately represent all possible classes that exist in the investigated cases. The quality of data is also an important factor. Training an untrained CNN requires monumental computational effort to set its parameters correctly. This can take weeks to months of calculation on a large-scale cluster of servers. CNN’s require a large number of images to achieve high prediction accuracy, for example, ImageNet contains over 14 million images in 21841 synsets [4].

Medical image databases are typically much smaller, with limited annotation of each image. They also tend to be restricted to specific studies of diagnostic interest. Data availability is also restricted due to privacy concerns and sharing policies of the acquiring medical centers. Fortunately, it is now possible to utilize transfer learning to interpret medical images. This technique allows the use of networks previously well-trained on millions of everyday images, and retrain them to classify medical images based on previously learned image features like shape, contours, intensity gradients and adjusting them to recognize characteristic features in medical images. This is still an emerging field, with only a few recent reports of transfer-learning CNN’s exhibiting satisfactory accuracy for medical images [5,6]. These include the classification of retinal images and diagnosis of skin conditions. The accuracy of such CNNs reaches ∼ 95%, generally considered to be adequate, and comparable to human interpretation of the images.

A very recent study [7] utilized transfer learning to detect adult wrist fractures in radiographs, but to our knowledge, no previous studies have successfully applied transfer learning to the problem of fracture detection in *pediatric* patients which are more complicated given the presence of growth plates and developing epiphyseal ossification centers that changes based on age. The purpose of this study is to apply transfer learning for AI augmented binomial detection of pediatric tibial fractures using a limited training set.

## Methods

### Dataset selection

This work was conducted under a protocol approved by the Institutional Review Board (IRB). We retrieved retrospective 516 studies of the foot, ankle, tibia and fibula conducted at a tertiary care pediatric hospital in the period 2009-2017. Mean age was 6.4±4.4 years, male to female ratio was 1.2. The studies contained 2118 radiographs, i.e., 2-3 images per study in multiple varying orthogonal views. Studies were classified, based on the official radiology report generated by Certificate of Added Qualification (CAQ) radiologist, into 3 groups: (1) normal (2) fracture (3) combined group: images containing fixation, patient in cast and other pathology. In this work, the third group was excluded since it did not meet the target of acute tibial fractures. From the remaining cases, we selected 490 studies fulfilling selection criteria (class 1 & 2), containing equal numbers of normal and fracture exams. The determination of “fracture” was inclusive of any transverse, oblique, longitudinal, Salter-Harris, buckle, comminuted, angulated, or displaced tibial fractures. Plastic bowing fractures without a discrete fracture line were excluded.

The radiographic fracture description of the 49 positive studies (19 ap view, 10 lateral, 20 oblique view) in the test set are detailed in Table 1.

**Table 1.** Radiographic fractures’ description of the test set

| Fracture type | Buckle | Oblique | Salter-Harris | Comminuted | Healing |
| --- | --- | --- | --- | --- | --- |
| **Displaced** = 15 | 1 | 10 | 3 | 1 | 0 |
| **Nondisplaced** = 34 | 6 | 22 | 6 | 0 | 7 |
| **Total** = 49 | 7 | 32 | 9 | 1 | 7 |

### Preprocessing

The preprocessing routine is shown in Figure 1. Radiographs were downloaded in DICOM format to a secure workstation, and de-identified. Each image was rotated to a standard position (Figure 1b), cropped to include the distal tibia, distal fibula and ankle joint and part of the talar dome (Figure 1c). All indexing letters were removed by masking with an empty square. Images were transformed to the size acceptable by the pre-trained network and normalized to (0-255) levels, and saved in 16-bit png format (Figure 1d). The images were then randomly divided into a training set/validation-set/test-set in an approximately 80/10/10 ratio: The training set contained 784 images, while the Validation set and the Test set each contained 98 images. All sets contained equal numbers of normal and fracture radiographs. Illustrative processed images are shown on Figure 2.

**Figure 1).**
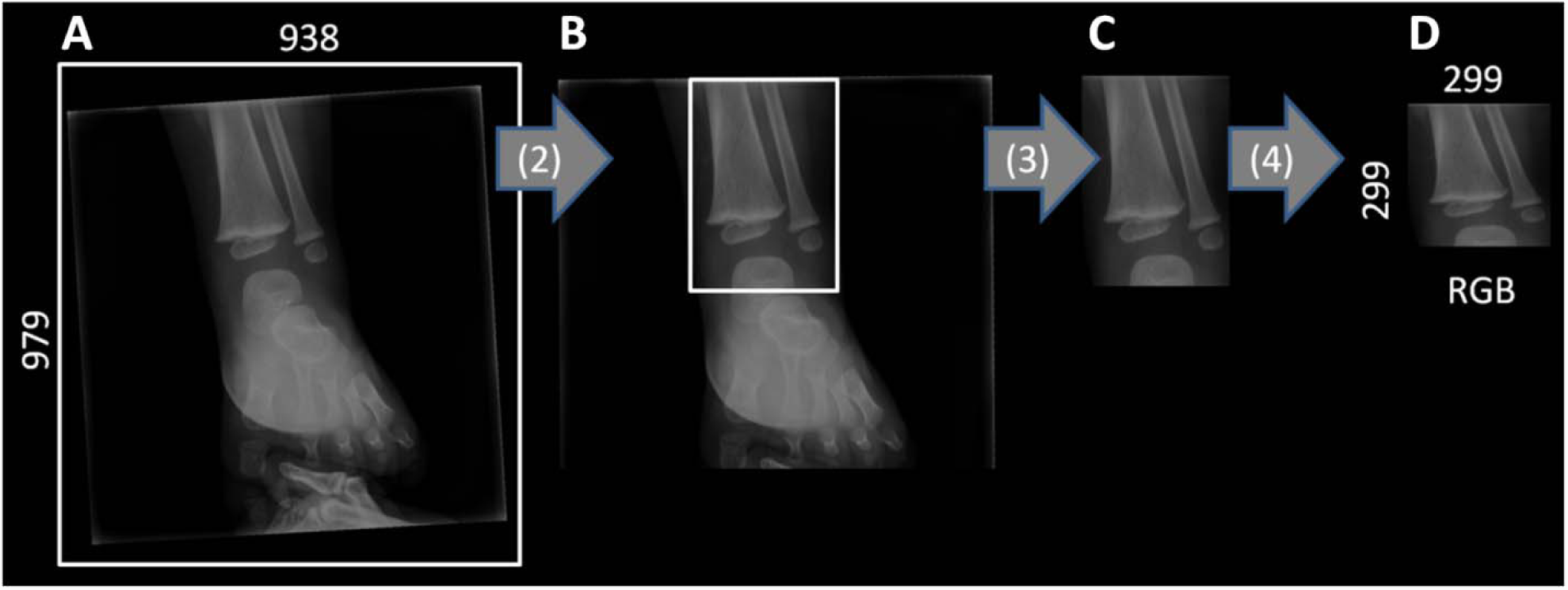
Preprocessing images for CNN input. (1) Anonymized radiographs, 2) Rotation to standard position, 3) cropping to ROI, 4) converting to square 299×299 RGB image in 16-bit png format.

**Figure 2).**
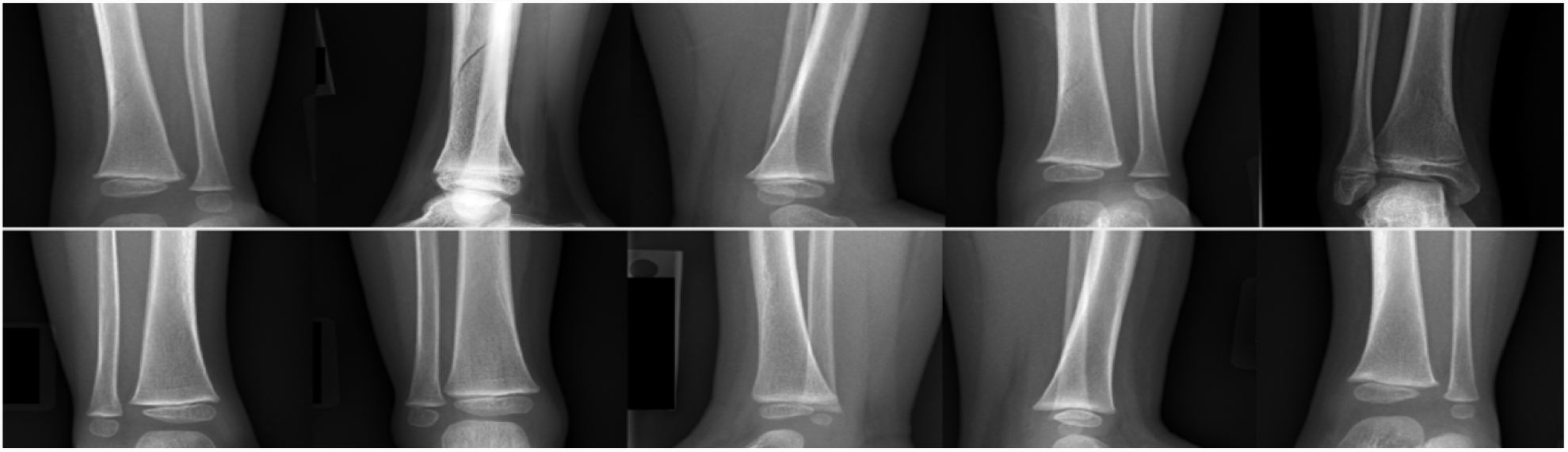
Example negative and positive representative images for toddler Tibial fractures for CNN network.

### Network architecture

We used a transfer learning approach based on the Xception architecture [8]. The Xception network was created on the base of the well know Inception v3 network, weights of the Xception network were obtained via training on 60 NVIDIA K80 GPUs within 72 hours of training. It outperforms Inception v3 in the classification of 17000 classes of images. For our purposes, the following modifications were made to the Xception network: 2 modules of fully convoluted reasoning layers with 5×128 nodes each, and incorporating drop-out segments were placed on top of the existing network. Drop-out segments prevent overfitting of the model to training data, and allow preferential matching of robust features correlated to fractures detection [9]. The Xception architecture consists of 36 convolutional layers structured into 14 modules, creating a feature recognition foundation of the network. In our transfer learning approach, we conserve weights of the first 9 modules allowing for training of the next 5 Xception modules plus 5 reasoning additional modules not included in the original Xception network.

### Network training

Network implementation and training were conducted in the Keras 2.1.4 environment with Python 2.7.12, and Tensorflow 1.3.0, on one Nvidia Tesla K80 GPU, on a dual Xeon CPU workstation.

### Data augmentation

Images in the training dataset were used to create an expanded training set using the following augmentations: (1) image feature normalization process which the subtraction from each image the average intensity of the group and dividing by its standard deviation (2) width and height shifts up to 5% (3) 5% zoom and (4) up to 20 degrees rotation (5) and horizontal flip. Augmentation process creates 8 images based on each input image. Using randomly assigned augmentations parameters augmentation expands overall input training set, and improve the resulting network accuracy.

### Training

The training was done in two phases. In phase 1, a short training set preliminary weights in the top 2 reasoning layers over 10 epochs was used. In phase 2, we adjusted the weights of pre-trained 5 layers of Xception and the 2 additional reasoning layers built on the top of the existing network to fine-tune the best accuracy in the validation set. Phase 2 was done over 300 epochs with the Nesterov Adam optimizer (NADAM) [10,11]. The final training period took ∼230 minutes.

## Results

The test set contained 98 images which were not available to the network during training and validation phases. The distribution of classification results as a function of model output is shown as a bar plot in figure 3. True positive (TP) and true negative (TN) cases are color coded. The trained network exhibits accuracy of 97.9% for accuracy, (95.9% sensitivity and 100% specificity), these results are summarized in Table 2. The receiver operating characteristic (ROC) curve illustrates the very high diagnostic ability of this binary classifier (Figure 4), with an area under the curve (AUC) equal to 0.995.

**Figure 3).**
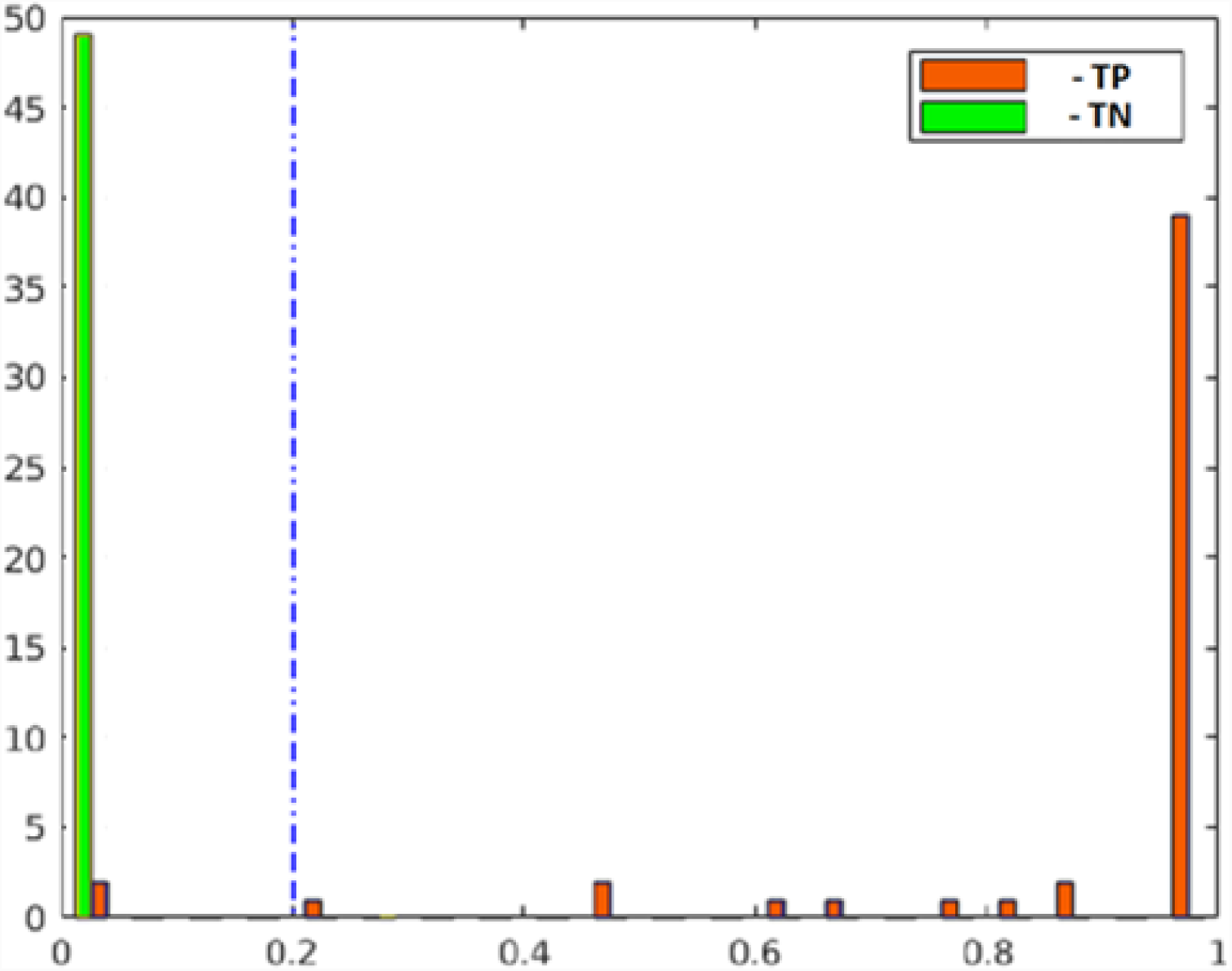
Bar plot showing distribution of classification results for the acute fracture (True positive) in red and the normal images-(true negative) in green form 100 test images. A higher score indicates a higher probability of a fracture. Dotted blue line class indicated chosen separation threshold TH = 0.2.

**Table 2.** Sensitivity, specificity, PPV, NPV, and accuracy of Transfer learning CNN radiographic evaluation of pediatric acute tibial fractures, for separation threshold = 0.2

|  |  |  |  |
| --- | --- | --- | --- |
| **CNN:**<br>**positive** | **47**<br>(A-True Positive) | **0**<br>(B-False Positive) | **PPV=100%**<br>$A/(A+B)*100\%$ |
| **CNN:**<br>**negative** | **2**<br>(C-False Negative) | **49**<br>(D-True Negative) | **NPV=96.1%**<br>$D/(D+C)*100\%$ |
| | **Sensitivity= 95.8%**<br>$A/(A+C)*100\%$ | **Specificity=100%**<br>$D/(D+B)*100\%$ | **Accuracy=97.9%**<br>$(A+D)/(A+B+C+D) *100\%$ |

**Figure 4).**
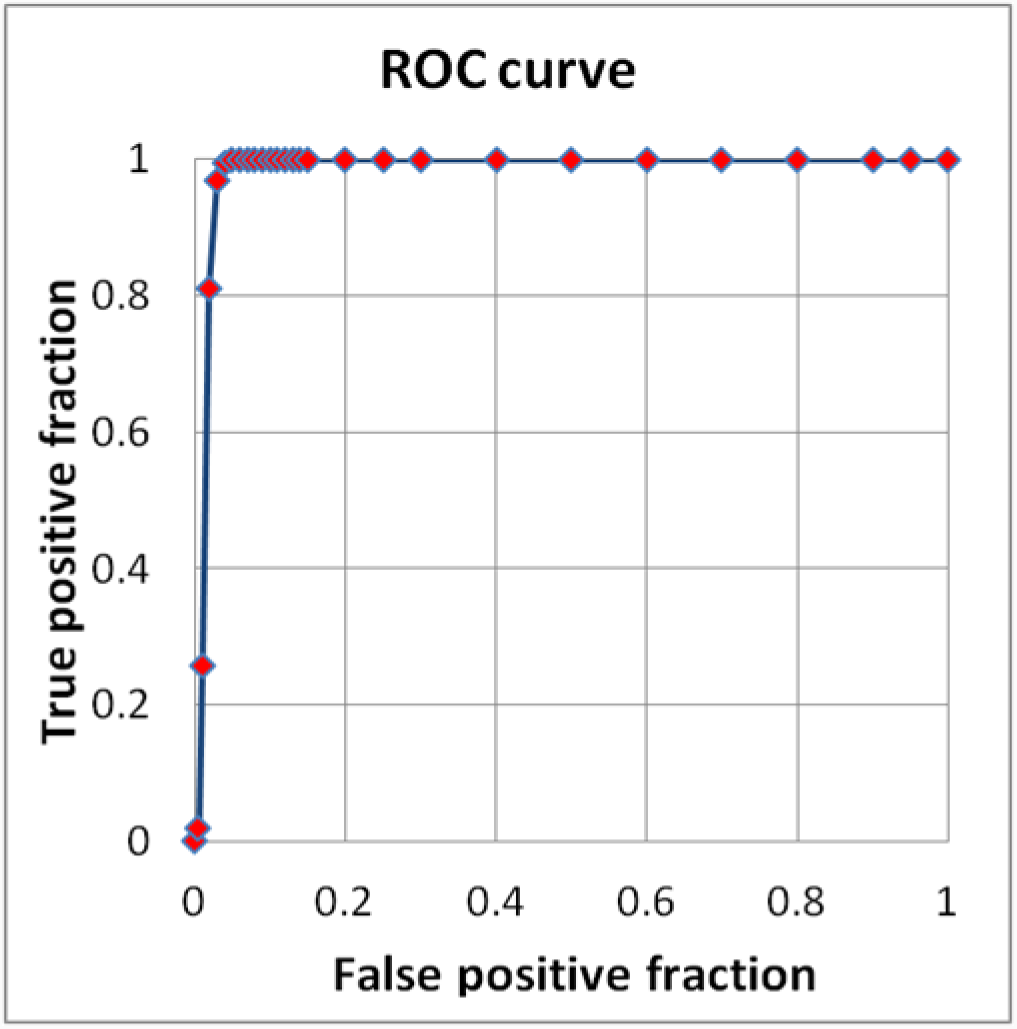
ROC curve. <u>RED symbols and BLUE line</u>: Fitted ROC curve. Fitted ROC Area: 0.995.

Maps showing areas for most activated neurons (Heat maps) are presented in figure 5, the top row corresponds to images with fractures, and shows that network activation correlates with the location of fractures. The bottom row show how trained network processed normal exams, where network activation correlates The most computation efforts correlates in regions outside the tibia and fibula, suggesting that the information in the fibula and tibia was processed without finding characteristic features of fractures, and as a result the network focused on the borders that contained edges. In the test set of 98 images, of 49 normal exams, none were mis-classified while 2 of 49 exams with fracture were classified as normal. These two images are shown in Figure 6, the nondisplaced oblique fracture line is not visible on figure 6A, while the case in figure 6B is a very poor quality, noisy image. It is not surprising that these two cases were wrongly read as normal.

**Figure 5).**
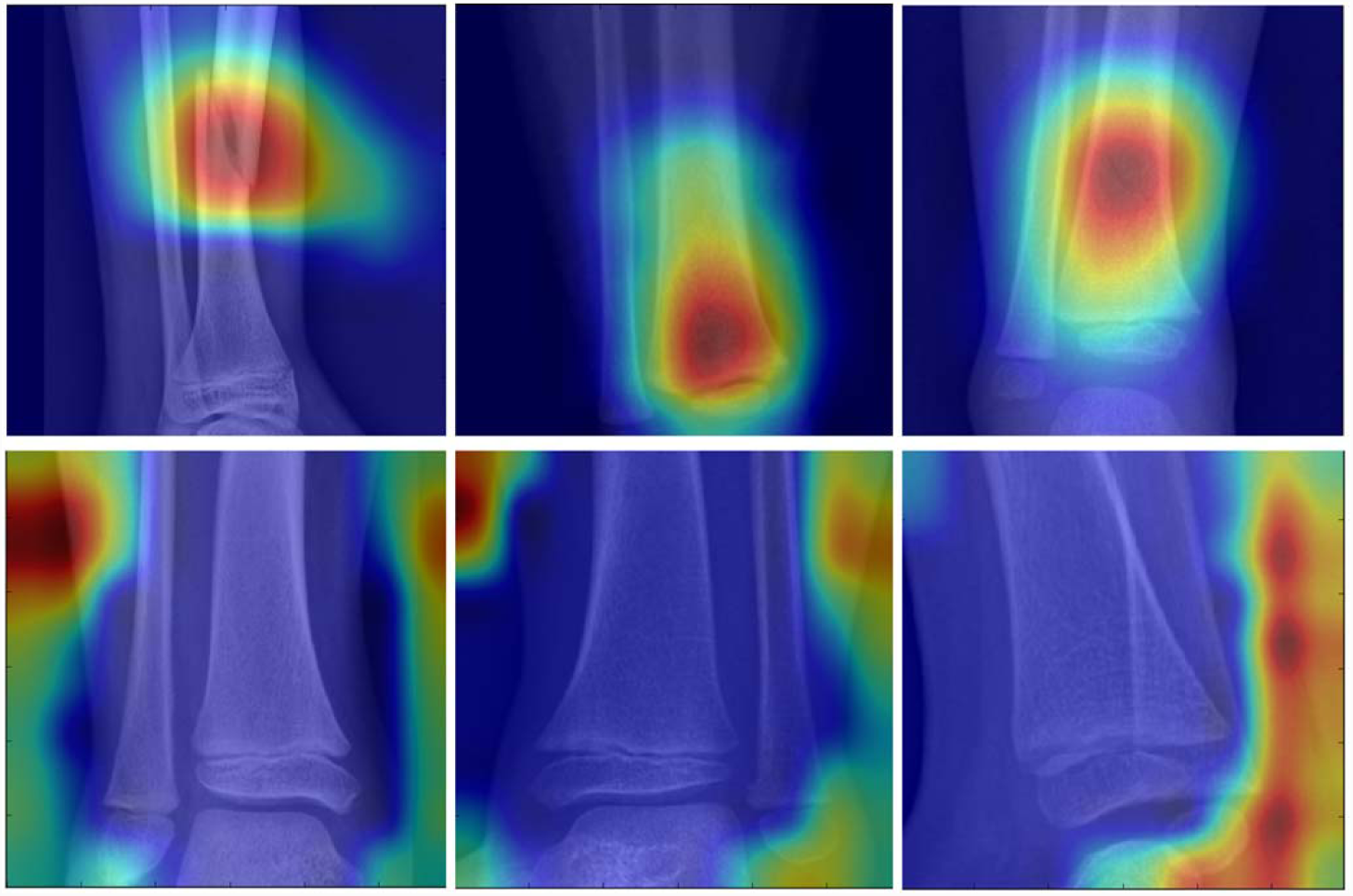
Heat maps example (color scale: red - high, blue - low) showing maps for activated neurons overlaid with investigated images, top row - positive images of fracture, bottom row negative - normal exams.

**Figure 6).**
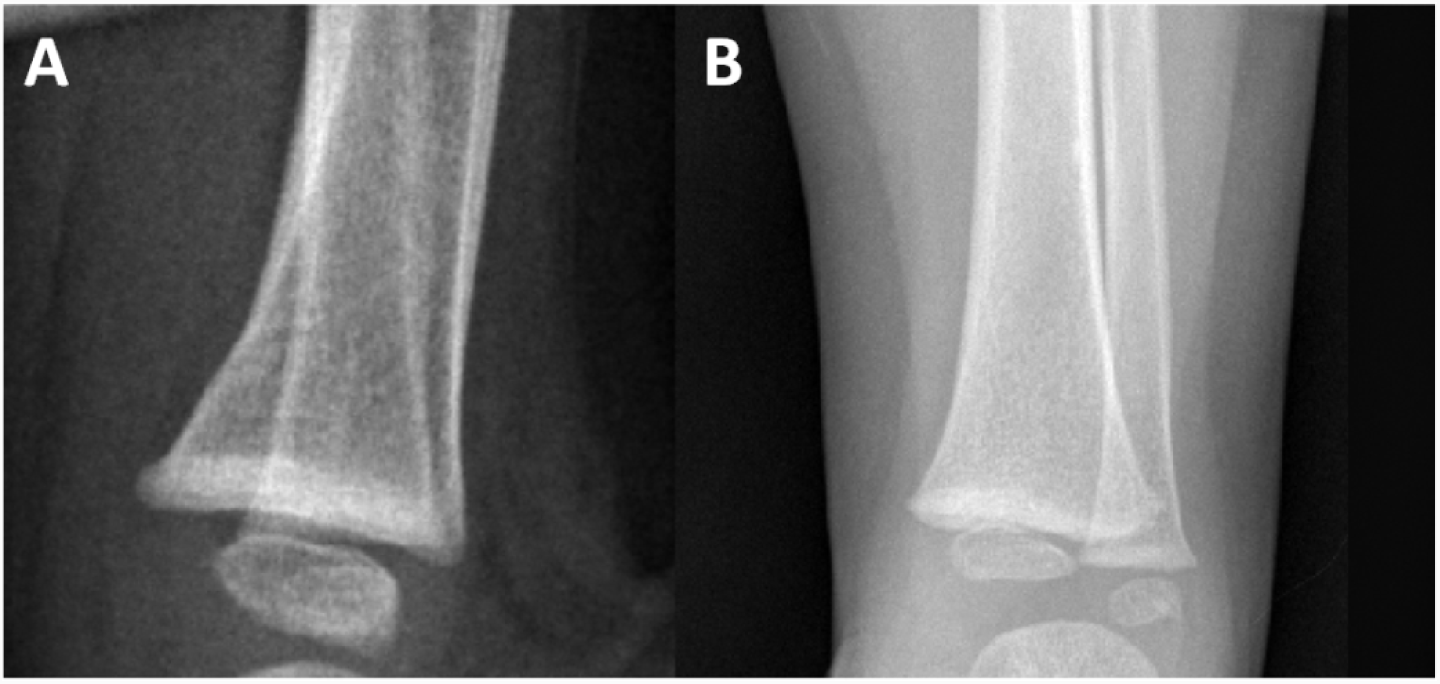
Two false negative (FN) images classification by CNN on the test set.

## Discussion

Recognizing pediatric tibial fractures is unique and distinct from adult tibial fractures due to the heterogeneous morphology related to varying sizes of growth plates, epiphyseal secondary ossification centers, and the variations in corticomedullary thickness predicated on age, nutrition, and body habitus of the child. As a consequence, there are many variations of pediatric long bone fractures that are unique and not seen in the adult population. A subspecialty trained pediatric radiologist is often available at a tertiary care center but is rarely accessible at community ER’s and trauma centers. Yet, subspecialty training is critical for accurate reads [12-15]. Capturing the essence of subspecialty trained radiologist reads in a computer algorithm that could be disseminated widely would therefore be highly beneficial. The success of our approach points the way to such algorithmic assistance tools that can be used by general radiologists, when a subspecialty trained pediatric MSK radiologist is not accessible.

Numerous CNN’s for diagnosing a range of pathologies have emerged in recent years. However, one uniform feature has been the need for large training datasets to adequately train the network for accurate performance. One approach to reduce training set requirements is transfer learning, which allows the utilization of a network previously trained on millions of images to be adapted to work with specific tasks. Thus, a network previously trained to recognize broad classes of images can be re-trained for the specific classification task of recognizing hairline fractures in X-ray images. While transfer-learning has played a role in reducing the training set requirements, the majority of networks still need tens of thousands of training cases to perform adequately. In this work we demonstrate that a CNN trained on as few as ∼900 images to optimize its weights, is capable of recognizing acute distal tibial fractures. The key to this accuracy is the preprocessing of images both in the training and test sets: restricting the anatomy to what is truly needed, fixing the image orientation, and maintaining uniform image size. These tasks can easily be automated, and do not constitute a significant burden.

This study is also the first to our knowledge that implemented a successful approach of providing a heterogeneous group of radiographs with different orthogonal projections to binomial fracture determination in skeletally immature patients, whereas other studies have only included a single orthogonal plane in adult patients [16,17]. Our results, despite the various different orthogonal views provided to the training and test set performed at a similar level of accuracy compared with similar and larger datasets using a single frontal view in adult patients for pediatric tibial fractures. Our statistical performance is on par to these aforementioned studies by Chung et al and Urakawa et al despite the fact that our studies included multiple different orthogonal planes and varying skeletal maturity of the tibia. We attribute our success to the transfer-learning approach to optimization of the network and training using a carefully curated training data set.

Growth plates and the fragmentary appearance of the epiphyseal ossification centers may superficially mimic the appearance of fractures but our CNN algorithm was trained successfully to differentiate these normal developmental findings from true pathology, inclusive of several Salter-Harris fractures. To our knowledge, this is the first study to report this observation.

A typical pediatric musculoskeletal radiologist, during his/her entire training (consisting of a 1 year internship, a 4 year residency in general radiology, a 1 year general pediatric radiology fellowship and a 1 year pediatric MSK fellowship), would likely see 700-1000 tibial fractures. It is interesting to note that our transfer learning approach with carefully pre-processed images, achieves high accuracy (∼99%) with a similar number of training set images. To our knowledge, this is the first time that such accuracies have been achieved using training sets that compare in size to those experienced by human practitioners.

## Limitations

This study was limited to images acquired at, and read at, a single institution. The network was trained and tested on acute fractures, and was not validated on images of partially healed fractures, images with any fixation in the field of view, or other pathology present. Positive (fracture) images chosen for the training set were single views that showed the fracture clearly. Input data was pre-processed, cropped, rotated and aligned thus exposing the network to very restricted images of the distal Tibia, Fibula and a portion of the talar dome with the centerline of the tibia aligned to the vertical direction. The input resolution was limited to 299×299 to accommodate transfer learning based on the Xception network. The reported results were obtained using single plane radiographs for the classification task in contrast to clinical settings where radiologists review multiple views before making a diagnosis.

## Conclusion

Our initial results in applying transfer learning demonstrates that CNN can robustly adapt and differentiate binomial variations of the normal skeletally immature tibia from fracture pathology with a limited training set with multiple different orthogonal views. This application of transfer learning for imaging interpretation may potentially be applied to other anatomic areas and other imaging modalities and is able to successfully differentiate acute pathology from normal variations of skeletally immature bones.

## References

[1] Lindsey R, Daluiski A, Chopra S, Lachapelle A, Mozer M, Sicular S, Hanel D, Gardner M, Gupta A, Hotchkiss R, Potter H. Deep neural network improves fracture detection by clinicians. Proc Natl Acad Sci U S A. 2018 Nov 6;115(45):11591–11596. doi:10.1073/pnas.1806905115

[2] Anu TC., Mallikarjunaswamy MS, Raman, R. Detection of bone fracture using image processing methods. IJCA proceedings on national conference on power systems and industrial automation NCPSIA. vol. 3.; 2015: 6–9

[3] Schmidhuber J. Deep learning in neural networks: An overview. Neural Networks 61 (2015) 85–117

[4] Russakovsky O,Deng J, Su H, Krause J, Satheesh S, Ma S, Huang Z, Karpathy A, Khosla, Bernstein M, Berg AC, Fei-Fei L. ImageNet Large Scale Visual Recognition Challenge.(2015) arXiv:1409.0575

[5] Shin HC, Roth HR, Gao M, Lu L, Xu Z, Nogues I, Yao J, Mollura D, Summers RM. Deep Convolutional Neural Networks for Computer-Aided Detection: CNN Architectures, Dataset Characteristics and Transfer Learning. IEEE Trans Med Imaging. 2016 May;35(5):1285–98. doi:10.1109/TMI.2016.2528162

[6] Choi JY, Yoo TK, Seo JG, Kwak J, Um TT, Rim TH (2017) Multi-categorical deep learning neural network to classify retinal images: A pilot study employing small database. PLoS ONE 12(11): e0187336. doi.org/10.1371/journal.pone.0187336

[7] Kim DH, MacKinnon T. Artificial intelligence in fracture detection: transfer learning from deep convolutional neural networks. Clinical Radiology 73 (2018) 439e445

[8] Chollet F. Xception: Deep Learning with Depthwise Separable Convolutions. arXiv:1610.02357

[9] Hinton G, Krizhevsky A, Sutskever I. et al. Dropout: a simple way to prevent neural networks from overfitting. J Mach Learn Res. 2014; 15: 1929–1958

[10] Dozat T. Incorporating Nesterov Momentum into Adam. ICLR Workshop, (1):2013–2016, 2016

[11] Rudner S. An overview of gradient descent optimization algorithms. arXiv:1609.04747v2

[12] Bisset GS 3rd, Crowe J. Diagnostic errors in interpretation of pediatric musculoskeletal radiographs at common injury sites. Pediatr Radiol. 2014 May;44(5):552–7. doi:10.1007/s00247-013-2869-9

[13] Eakins C, Ellis WD, Pruthi S, Johnson DP, Hernanz-Schulman M, Yu C, Kan JH. Second opinion interpretations by specialty radiologists at a pediatric hospital: rate of disagreement and clinical implications. AJR Am J Roentgenol. 2012 Oct;199(4):916–20

[14] Kung JW1, Melenevsky Y, Hochman MG, Didolkar MM, Yablon CM, Eisenberg RL, Wu JS.On-call musculoskeletal radiographs: discrepancy rates between radiology residents and musculoskeletal radiologists. AJR Am J Roentgenol. 2013 Apr;200(4):856–9. doi:10.2214/AJR.12.9100

[15] Tomich J, Retrouvey M, Shaves S. Emergency imaging discrepancy rates at a level 1 trauma center: identifying the most common on-call resident “misses”. Emerg Radiol. 2013 Dec;20(6):499–505. doi:10.1007/s10140-013-1146-4

[16] Chung SW, Han SS, Lee JW, Oh KS, Kim NR, Yoon JP, Kim JY, Moon SH, Kwon J, Lee HJ, Noh YM, Kim Y, Automated detection and classification of the proximal humerus fracture by using deep learning algorithm. Acta Orthop 2018 89(4):468–473, doi:10.1080/17453674.2018.1453714

[17] Urakawa T, Tanaka Y, Goto S, Matsuzawa H, Watanabe K, Endo N. Detecting intertrochanteric hip fractures with orthopedist-level accuracy using a deep convolutional neural network. Skeletal Radiol. 2018 Jun 28. doi:10.1007/s00256-018-3016-3

